# PeptideForest: Semisupervised machine learning integrating multiple search engines for peptide identification

**DOI:** 10.1101/2022.12.21.521351

**Authors:** T. Ranff, M. Dennison, J. Bédorf, S. Schulze, N Zinn, M. Bantscheff, J.J.R.M. van Heugten, C. Fufezan

## Abstract

The first step in bottom-up proteomics is the assignment of measured fragmentation mass spectra to peptide sequences, also known as peptide spectrum matches. In recent years novel algorithms have pushed the assignment to new heights, unfortunately, different algorithms come with different strengths and weaknesses and choosing the appropriate algorithm poses a challenge for the user. Here we introduce PeptideForest, a semi-supervised machine learning approach that integrates the assignments of multiple algorithms to train a random forest classifier to elevate that issue. Additionally, PeptideForest increases the number of peptide-to-spectrum matches that exhibit a q-value lower than 1% by 25.2 ± 1.6% compared to MS-GF+ data on samples containing mixed HEK and *E. coli* proteomes. However, an increase in quantity does not necessarily reflect an increase in quality and this is why we devised a novel approach to determine the quality of the assigned spectra through TMT quantification of samples with known ground truths. Thereby, we could show that the increase in PSMs below 1% q-value does not come with a decrease in quantification quality and as such PeptideForest offers a possibility to gain deeper insights into bottom-up proteomics. PeptideForest has been integrated into our pipeline framework Ursgal and can therefore be combined with a wide array of algorithms.

## Introduction

Mass Spectrometry is altering proteomics at a high pace with more advanced instruments being developed every year with advancements in engineering and software development. A key step in bottom-up proteomics is the assignment of peptides with potential modifications to observed mass spectra, also known as peptide-spectrum matches (PSMs). Several approaches exist to assign PSMs, such as database search ^1–8^, spectral library search ^9–13^, *de novo* sequencing ^14–17^, and open modification search ^8,18–20^. The most common approach, the database search, generates theoretical spectra from a list of sequences and compares them to the measured spectra. Although database search is the most established approach, the fact that innovative algorithms are still being published in recent years ^8,21,22^ shows that the problem of assigning spectra to peptides with confidence has not been fully solved yet. Additionally, different PSMs assignment strategies implemented by different database search engines always bring certain benefits and drawbacks and thus it is difficult from a user perspective to choose the appropriate one for the question at hand. Furthermore, even the combination of results from different search engines is non-trivial since scoring algorithms differ. Some search engines like Sequest score a spectrum based on the assignable and non-assignable peaks without taking e.g. database size into account ^1,23^. Other engines like X!Tandem uses a probabilistic approach taking all possible assignments for one given spectrum into account ^24^.

Several approaches exist that combine different search engine results ^25–29^ yet so far none of them has employed machine learning to increase the overall number of PSMs and most importantly evaluate the quality of the increased number of PSMs at a given q-value. For single search runs, Käll et al. pioneered the application of machine learning with the release of Percolator ^30^, which helps to boost the number of PSMs using support vector machines by separating target and decoy assignments better. Percolator implements a three-fold cross-validation approach: the PSMs are split into three random sets on the spectrum level ^31^. During training, the model is fitted to two of the three sets, while predictions are made on the remaining set. This is repeated for each of the pairs of sets, such that each PSM is scored by a model that has not been trained using these PSMs.

Since machine learning algorithms depend on the features they are trained on, we decided to develop an approach that takes the results for multiple search engines into account and can be extended with ease. Our approach has been facilitated by Ursgal, a library wrapping multiple search engines in a unified framework ^29^. The advantages are twofold: a) each search engine can be employed using the same general parameter set since parameter translation is done by Ursgal and most importantly b) the output of each search engine is unified. For the latter, we have devised a standalone Python package called pyProtista (https://github.com/computational-ms/pyProtista), which will be presented elsewhere.

Here, we present a novel approach called PeptideForest, a random forest regression machine learning implementation to rerank PSMs based on the search results from multiple database search engines. Our results show an increase of 23-27% PSMs at 1% q-value threshold. Furthermore, we used a known truth based on isobaric mass tag-encoded tryptic digests of human and *E*.*coli* protein extracts to benchmark assignment quality. The additional PSMs matched the expected quantification profiles therefore demonstrating that the increased number of PSMs does not reduce the quality of the assignments.

## Results

We built on the approach of Percolator, first described in ^30^, which uses a Support Vector Machine (SVM) machine learning model to separate high-quality target PSMs (defined as those with a *q-value <= 1%*) from decoy PSMs. Our approach, PeptideForest, uses random forest regression (RF-reg) to predict whether a PSM is a target or a decoy, with the probability that a PSM is a target used as a score, which is used to re-rank PSMs and identify more high-quality target PSMs as matches than the search engines alone.

The performance of PeptideForest in comparison to different database search engines is shown in figure 1. The number of unique peptide sequences including their modifications, also known as peptidoforms, are plotted against different q-value thresholds (Fig. 1A) for Mascot (v2.6.2)^1^, OMSSA (2.1.9)^3^, MSFragger (v.3)^8^, MS-GF+ (v2021.03.22)^7^, Comet (2020.01.4)^38^, MSAmanda (2.0.0.17442)^39^, and PeptideForest (RF-reg, v.1.0) for the four test data sets of this study (E13, E32, E41, E50). The test samples contained a mix of human embryonic kidney (HEK) and *Escherichia coli* proteomes. On average, at 1% q-value, PeptideForest yields 20200 ± 1475 unique peptidoforms, while MS-GF+ as second best algorithm yields only 16144 ± 1378 unique peptidoforms (Fig. 1A). Over all test datasets, applying PeptideForest led to an increase in PSMs of 25.2 ± 1.6% compared to the best single search engine MS-GF+ (Fig. 1B). Percolator 3.5^30^, the current state of the art machine learning algorithm for single search results uses such multi-dimensional feature space and support vector machines. However, PeptideForest using multiple engines outperforms Percolator 3.5 using only MSFragger by 51 ± 8% (Fig. 1C) of additionally identified peptidoforms.

**Fig. 1:**
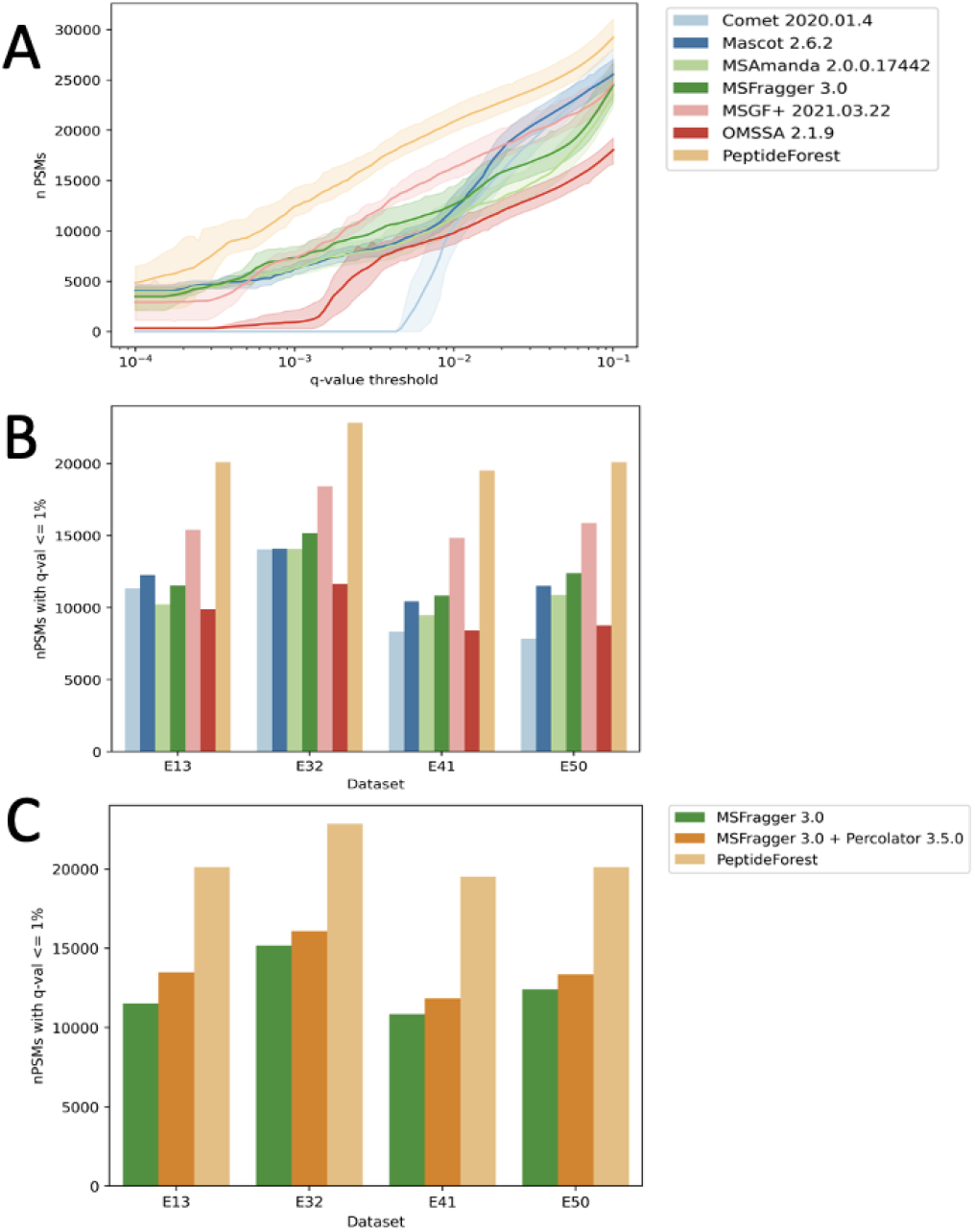
Comparison of search engines (indicated by legend) and PeptideForest. **(A)** Number of unique peptidoforms (y-axis) at different q-value thresholds (x-axis) across all datasets. Average shown as solid line, standard deviation as shaded surrounding area **(B)** Number of unique peptidoforms at 1% q-value for all data sets. **(C)** Number of unique peptidoforms (y-axis) identified by MSFragger, MSFragger + Percolator, and PeptideForest (indicated by legend) at a 1% q-value threshold across all datasets. MSFragger identified 12500 ± 1900, MSFragger with Percolator identified 13700 ± 1800, and PeptideForest identified 20700 ± 1500 peptidoforms respectively.

### Re-ranked PSMs

After showing that the number of peptidoforms identified above a given threshold could be substantially increased using PeptideForest, we further examined the basis of the underlying re-ranking. Figure 2 visualizes the q-values reported by all engines and compares it to PeptideForest in E13. Each row in the heatmap represents a PSM. The two major separations are a) PSMs where all engines agree on mostly a q-value <1% (clade C2 Fig. 2) and PSMs that show higher degree of variations (clade C1 Fig.2). In clade C1, two groups can be distinguished, clade C1A shows PSMs that were re-ranked by PeptideForest to be mostly below 1% although most engines evaluated q-values to be <5% and clade C1B where most engines report q-values to be above 5% yet PeptideForest still re-ranks most of those PSMs to be below 1%. For each of these categories four concrete PSMs (A - D, denoted on y-axis 2) were singled out and evaluated (see Figure 3).

**Fig. 2:**
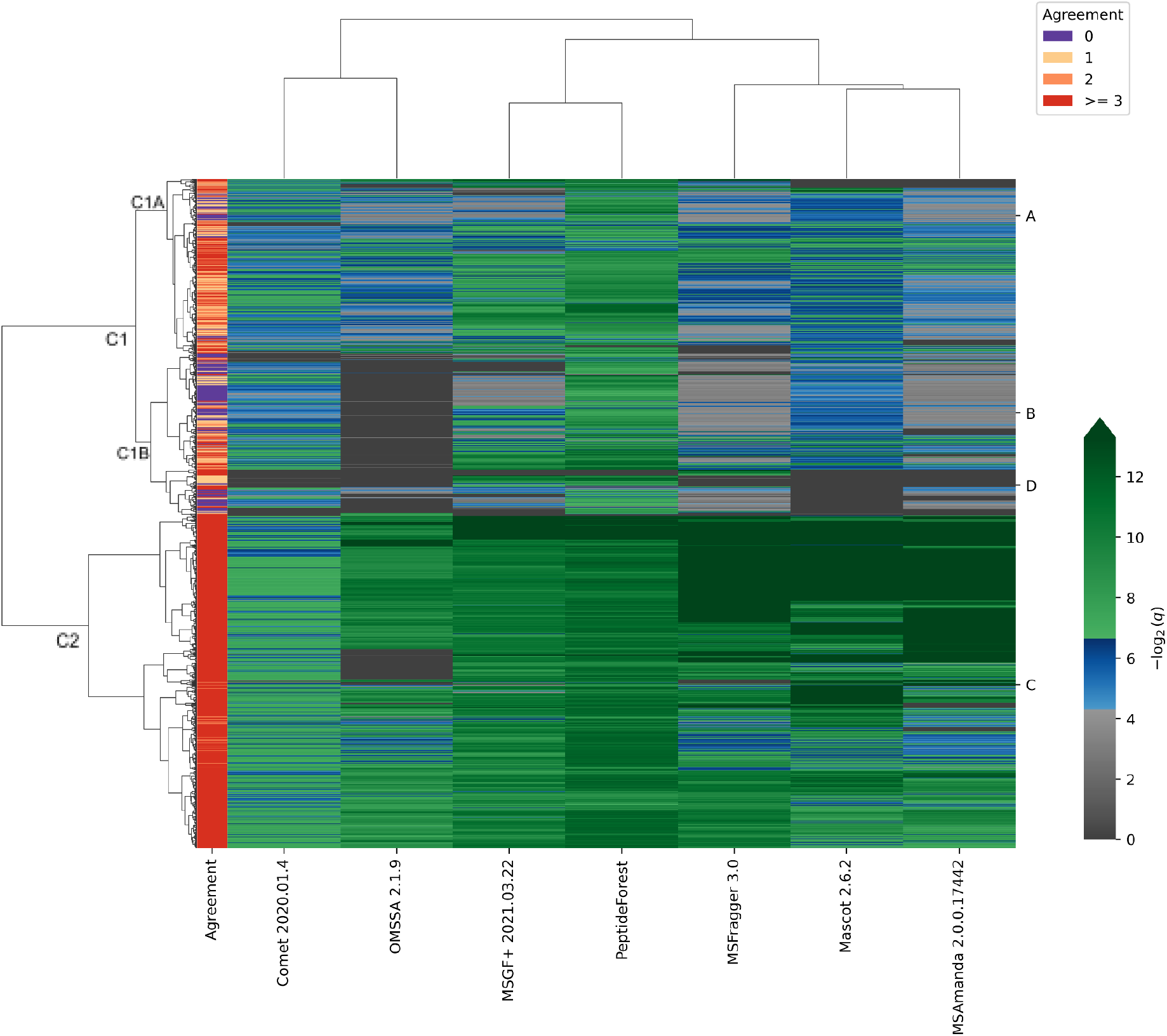
PSM q-values across all engines and agreement with PeptideForest in the E13 dataset where at least one q-value <1% was reported. Q-values from 0-1% are color-coded green, 1-5% blue, and 5-100% gray. Separately, the agreement between the engine scores and the PeptideForest score is evaluated (see legend). An agreement of zero (purple) meaning either no other engine assigned a q-value <1% when PeptideForest did or all engines assigned q-values of <1% when PeptideForest did not. If the agreement is ≥3 at least 3 engines ranked the PSM similarly to PeptideForest.

**Fig. 3:**
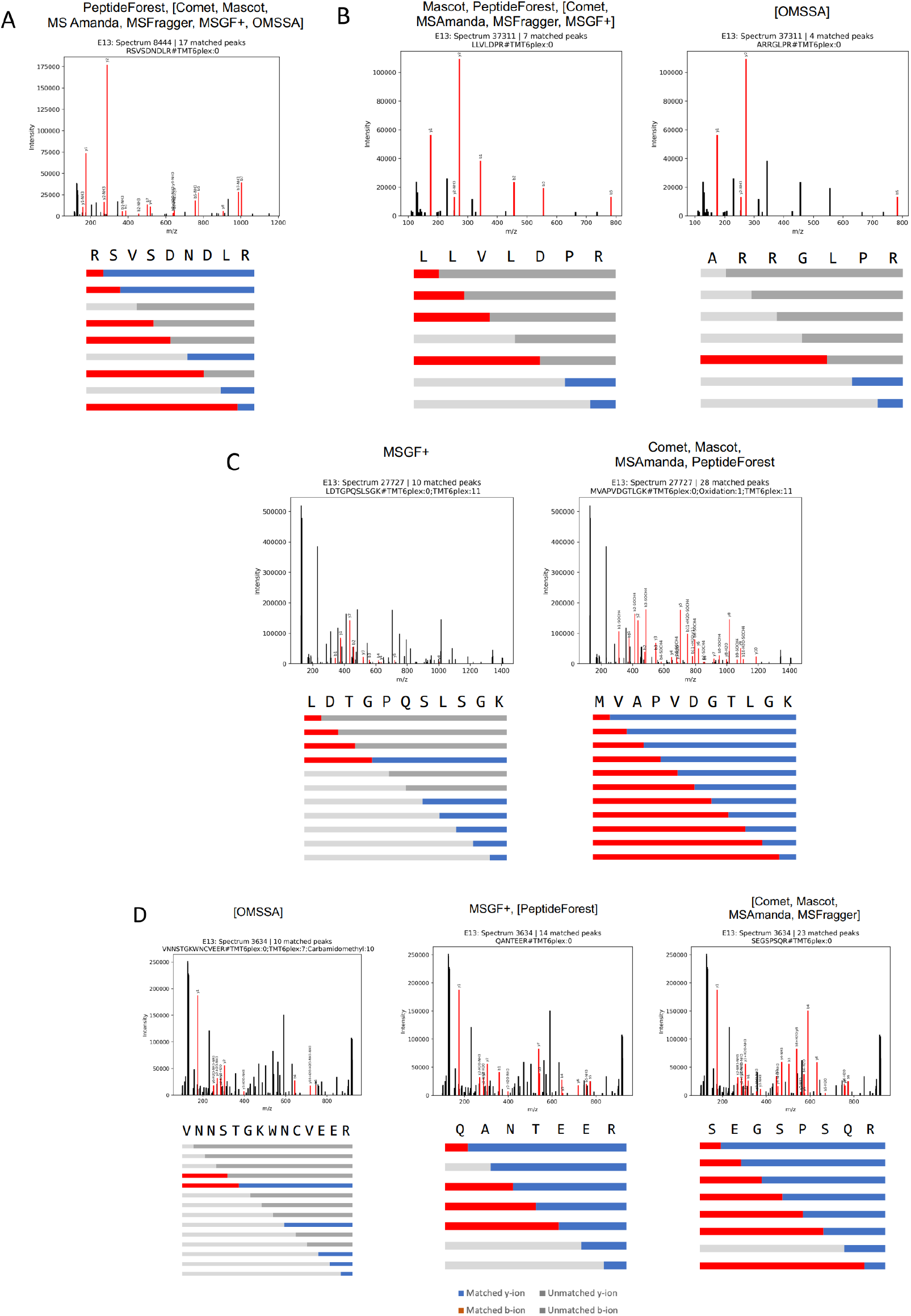
shows the examples highlighted in figure 2, search engine names are in brackets if PSM assignment is above 1% q-value, upper panel shows annotated MS2 spectra with matched ions, lower panel shows PSM base sequences and ion coverages (b and y ions as red and blue horizontal bars, respectively; missing ions are shown in gray, modifications are taken into account yet not explicitly listed). **(A)** RSVSDNDLR+mods, Spectrum ID 8444 - only PeptideForest assigns a q-value < 1%, **(B)** left panels LLVLDPR+mods found by all engines except OMSSA, but only Mascot and PeptideForest assigned a q-value < 1%, right panels ARRGLPR+mods the PSM assigned by OMSSA with q-value > 1%, **(C)** left panels LDTGPQSLSGK+mods assigned by MSGF+ with q-value < 1%, right panels MVAPVDGTLGK+mods by all other engines with q-value < 1%, **(D)** left panels VNNSTGKWNVEER+mods, only by OMSSA with q-value > 1%, middle panels QANTEER+mods by MSGF+ (<1%) and PeptideForest (>1%), right panel SEGSPSQR+mods, all other engines with q-value > 1%.

Figure 3 shows the examples of the categories described above and in Figure 2.

In Figure 3A, all search engines have reported the same PSM yet only PeptideForest pushed its q-value below 1%. The fragmentation spectrum (MS2) shows that nearly all peaks match the predicted PSM and the sequence coverage in the lower panel is supporting the re-ranking.

Figure 3B shows a similar case to 3A, all engines except OMSSA report the same PSM, yet only Mascot and PeptideForest would assign it with an q-value below 1 %. The MS2 spectrum shows that nearly all peaks have matched theoretical ions for the assigned PSM and although the sequence coverage is sparse the assignment is supported by the remaining four engines, even though they have not reported this PSM as a top hit (< 1%).

Another example is shown in Figure 3C, where all engines reported a PSM with a q-value < 1%, yet there was no agreement on the identity. MSGF+ reported LDTGPQSLSGK+mods (left panels) while all other engines report MVAPVDGTLGK+mods (right panels). The sequence coverage and matched ions clearly show that the assignment by MSGF+ is of lower quality compared to the other engines, i.e. the right panel, highlighting the correct decision made by PeptideForest.

Lastly, in Figure 3D PeptideForest re-ranks the assigned PSMs to be above the 1% threshold as the search engines show a high degree of ambiguity on identity level and on q-value category. MSGF+ is the only engine that would assign a q-value below 1% to QANTEER+mods whereas all other engines (except OMSSA) are reporting SEGSPSQR+mods but with a q-value above 1%. Arguably the latter PSM is better supported by the matched ions and sequence coverage and thus the re-ranking by PeptideForest is introducing a higher quality filter criterion, where one-engine wonders are penalized.

### Increasing quantity while maintaining quality

Commonly the number of PSMs or the number of unique peptidoforms within a given q-value cutoff is used as a measure to compare different algorithms, similar to Figure 1A. However, the legitimate question is whether or not the additional PSMs are actually of the same quality as the rest of the data and this is rarely addressed. Here we introduce consistency in relative quantification as a metric to assess the quality of PSMs. We mixed *E. coli* and human proteomes in well-defined ratios and labeled the different mixtures using isobaric mass tag reagents (TMT11) (Table 1). As a result, the assignment of a spectrum to *E. coli* or human can be done by using the quantification profile of the reporter ion without the need for identification. Equally, identifications can be validated by inspecting the reporter ion intensities. For example, given our mixing schema (Table 1), a peptide originating from an *E. coli* protein has the log2 ratio between reporter ion intensity 127L and 130L of log2(1/0.1), i.e. around 3.3. However, if a peptide exists in a human and an *E. coli* protein, then this log2 ratio will be a mixture of 3.3 and 0 weighted by the abundance of that peptide in the respective proteome. Equally, all other TMT channels that have a mixture of human and *E. coli* proteomes can be used for this kind of evaluation. Peptides existing purely in the human proteome will exhibit a log2 ratio of 0 for all combinations of channels excluding 127H, 128H, 129H, and 130H, which should be empty. Therefore, if we inspect all *E. coli* PSMs of a high-quality data assignment, we would expect the distribution of the log2 ratio between reporter ion intensity 127L and 130L to be around 3.3 and with a decrease in quality, we would expect to see more and more human peptides to be misassigned as *E. coli* thereby driving the log2 ratio distribution towards 0.

**Table 1:** Proteome mixtures at the different TMT channels.

| TMT | 126 | 127L | 127H | 128L | 128H | 129L | 129H | 130L | 130H | 131L |
| --- | --- | --- | --- | --- | --- | --- | --- | --- | --- | --- |
| *E. coli* | 0 | 1 | 1 | 0.5 | 0.5 | 0.2 | 0.2 | 0.1 | 0.1 | 1 |
| Human | 1 | 1 | 0 | 1 | 0 | 1 | 0 | 1 | 0 | 1 |

It is worth noting that the mixtures come with pipetting errors, ratio compression ^32^, and additional background isolation ^33^. Although we did correct for the latter two effects by filtering for S2I correction > 0.7, pipetting error was still an issue. Therefore, we are not using the expected ratio but we are rather using the differences in log2 ratio distributions to compare the different identification approaches regarding their assignments to *E. coli* peptides. As a result, we rely less on the accurate knowledge of the ground truth, yet we can estimate whether or not the distribution of the log2 ratios stems from the same original distribution. For example, if an increase in the number of PSMs comes with a decrease in quality, we would expect the overall distribution of proteotypic *E. coli* peptides to shift away from a log2 ratio defined by the *E. coli* mixture towards a log2 ratio defined by the human proteome background, that is towards a ratio of 0. As a result, the ratio distribution could be identified as not coming from the same original distribution using the Kolgomorov-Smirnov test.

As reported ions with higher intensities result in higher quantification accuracy and precision, we binned the data into 10 intensity bins defined by the maximum of the two reporter ion intensities investigated. As expected and shown in Fig. 4, the higher intensity bins reflect the expected log2 ratio of 3.3 for 127L and 130L better than the lower intensity bins. Importantly, the violin plots show similar distributions for Mascot, MSGF+, and PeptideForest in the respective bins. When applying the Kolgomorov-Smirnov test to compare engine distributions for each bin, only intensity bin 7 comparing Mascot to PeptideForest shows a p-value < 0.01, thus would indicate that the null hypothesis is accepted, i.e. that both distributions were sampled from different original distributions. For all other comparisons the null hypothesis cannot be rejected and thus the distributions are not significantly different, showing that the increase in quantity using PeptideForest does not come with a decrease in quality.

**Fig. 4:**
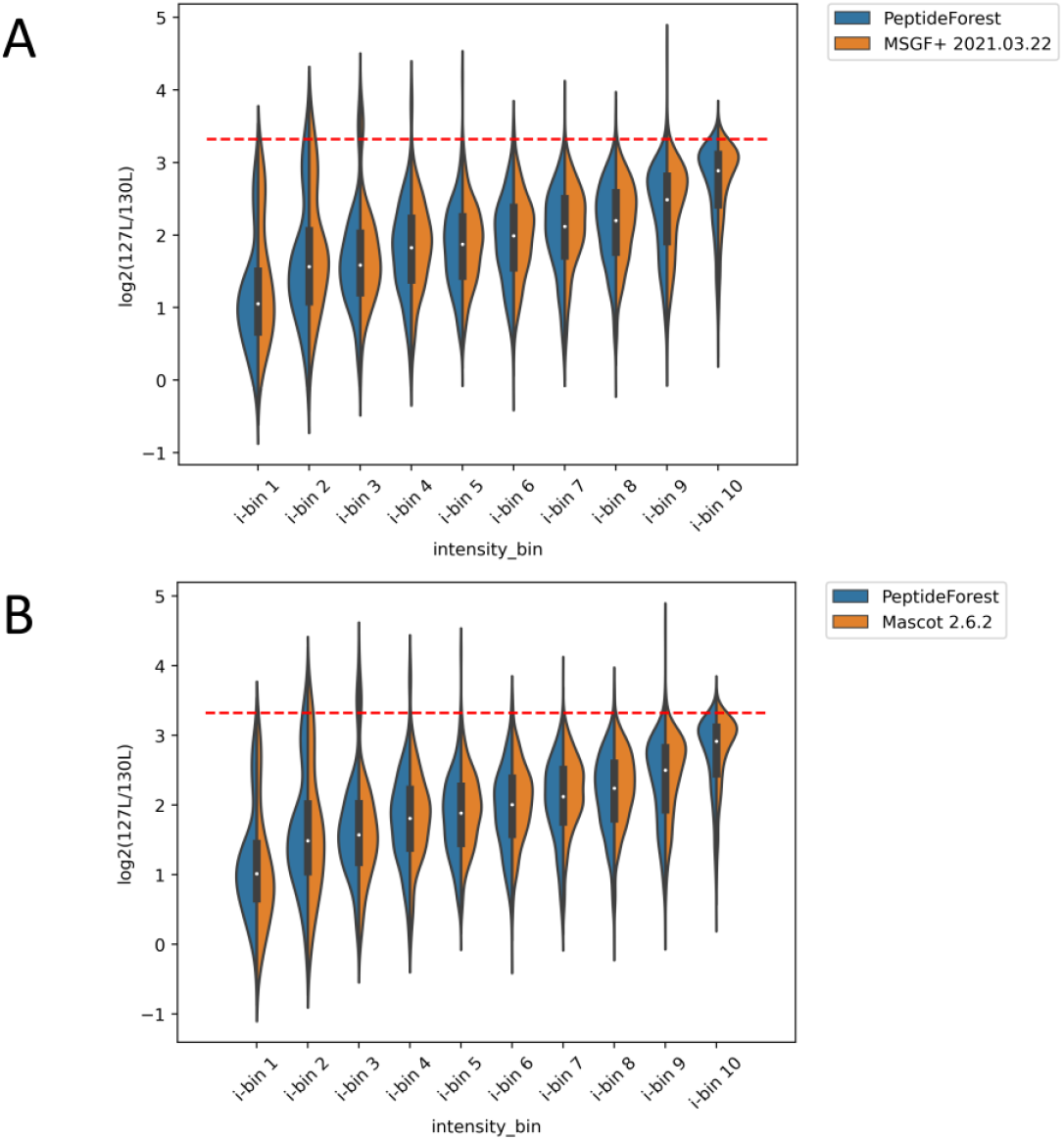
Violin plots showing the distribution of the log2 ratios 127L/130L for MSGF+ **(A)** and Mascot **(B)** (orange) versus Peptide Forest (blue) for ten intensity bins, i-bin0 to i-bin9, with i-bin 0 denoting the lowest intensity bin. High-intensity bins reflect the expected ratio of 3.3 better than low-intensity bins.

Finally, we tested all intensity bins for all log2 ratios employing the Kolgomorov-Smirnov test for all engine combinations. After correction for multiple testing using the method of Benjamini-Hochberg, few of the comparisons showed a p-value < 0.01 supporting the notion that the additional PSMs below 1% q-value cutoff resulting from processing with PeptideForest are not significantly different in quality than the single search engine results (Fig. 5). It is worth highlighting that those comparisons are on low to medium intensity bins and not systematically across all channels.

**Fig. 5:**
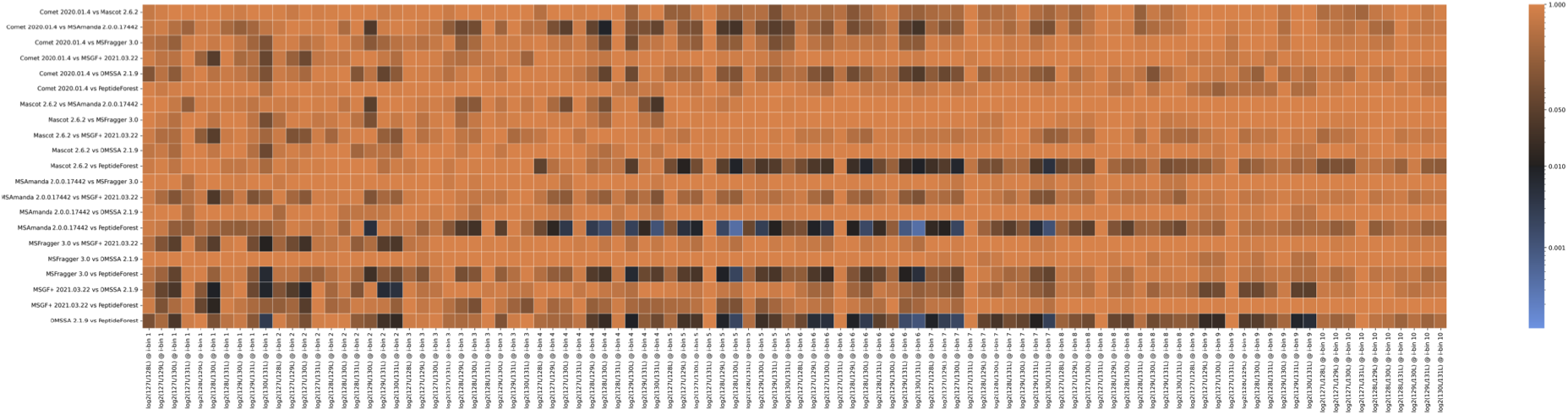
Kolgomorov-Smirnov test results comparing all engines against each other for all TMT channels that allow fold changes to be compared against the ground truth. Most of the comparisons are not statistically significant (color orange to black).

## Discussion

Here we introduce a semi-supervised machine learning approach employing random forest regression which we call PeptideForest. PeptideForest can re-rank target PSMs that resemble high-quality matches by using the scores of multiple search engines and other features. As a result, PeptideForest identifies 25.2 ± 1.6% more peptidoforms with a q-value < 1%, while maintaining the quality of identifications, as shown by the subsequent orthogonal validation approach that uses quantification profiles. The easiest underlying rationale behind the increase in peptide-to-spectrum matches (PSMs) is that PSMs assigned by multiple independent search engines representing consensus bear more weight. However, PeptideForest also introduces high quality filtering by re-ranking PSMs to be above 1% q-value when the individual engines have a high degree of variation, even if some report q-values of below 1%. This highlights the power of multiple data modalities, i.e. multiple engines using machine learning with a multi-dimensional feature space instead of merely using one engine.

Nevertheless, an increased number of PSMs may cause a drop in overall data quality. Our experimental setup allowed falsely assigned *E. coli* peptides to be identified based on their TMT quantification channels. The global analysis of quantification profiles of PSMs that were assigned to *E. coli* did not show any significant difference between PeptideForest and the other search engines, not even when compared to the most conservative engine in this study, Mascot 2.6.2. This supports the notion that the increased number of PSMs reported by PeptideForest does not reduce the quality of the data.

Here we showed that PeptideForest is a machine-learning algorithm that can help to push the boundaries of bottom-up proteomics by increasing the number of high-quality PSMs. PeptideForest is available on GitHub, is licensed under MIT, and is wrapped in Ursgal for convenience. Especially the integration into Ursgal makes the usage of multiple search engines very straightforward. Finally, PeptideForest has been designed in such a general way that it could easily be extended with new features such as results from open modification or *de novo* searches, also available in Ursgal. The combination of the different search types could also help to find appropriate cut-offs for *de novo* or other approaches that cannot employ target-decoy-based assessments.

## Material & Method

### Sample preparation

*E. coli* spiked into human background were produced by a tryptic digest of human and *E. coli* cell lysates as described in Smith et al. ^34^. In brief, proteins in 2% SDS were bound to Paramagnetic beads (SeraMag Speed beads, GE Healthcare, 45152105050250 and 651521050502) on filter plates (Multiscreen, Merck-Millipore, 10675743) by addition of ethanol to a final concentration of 50%. After four washes of the beads with 200 μl of 70% ethanol, beads were resuspended in trypsin and LysC in 0.1 mM HEPES, pH 8.5 containing TCEP and chloroacetamide and incubated at room temperature overnight. Peptides were pipetted at defined ratios according to Table 1 before labeling with TMT ^35,36^. The human background consisted of pooled tryptic digests from human cell lysates.

Peptides were labeled with isobaric mass tags (TMT10; Thermo Fisher Scientific) using the 10-plex TMT reagents, enabling relative quantification of ten conditions in a single experiment. The labeling reaction was performed in 100 mM HEPES, pH 8.5 at 22 °C and quenched with glycine. Labeled peptide extracts were combined into a single sample and lyophilized.

### Sample measurements

Samples were dried *in vacuo* and resuspended in 0.05% trifluoroacetic acid in water. Approximately 0.5 ug of peptide were injected into an Ultimate3000 nanoRLSC (Dionex) coupled to a Q-Exactive HF (Thermo Fisher Scientific). Peptides were separated on custom-made 50 cm × 100μm (ID) reversed-phase columns (1.9 μm C18, Reprosil) at 55 °C. Gradient elution was performed from 2% acetonitrile to 30% acetonitrile in 0.1% formic acid and 3.5% DMSO over 1 or 2h. The mass spectrometer was operated with a data-dependent top-ten method. Mass spectra were acquired using a resolution of 60,000 and an ion target of 3×10^6^. Higher energy collisional dissociation (HCD) scans were performed with 35% normalized collision energy (NCE) at 30,000 resolution (at an m/z of 200), and the ion target setting was set to 2×10^5^ to avoid ion coalescence^36^. The instrument was operated with Tune 2.5.0.2042 and Xcalibur 3.0.63.

### Mass spectra to peptide assignments

Mass spectrometry run files, RAWs were converted to mzMLs using msconvert^37^. MzML files were then searched using Mascot (v2.6.2)^1^, OMSSA (2.1.9)^3^, MSFragger (v.3)^8^, MS-GF+ (v2021.03.22)^7^, Comet (2020.01.4)^38^ and MSAmanda (2.0.0.17442)^39^ using carbamidomethylation of cysteine and TMT of lysine residues as fixed modification and oxidation of methionine, acetylation and TMT of protein N-termini as variable modifications. To compensate for machine drifts, the data was investigated using a wide search approach followed by a narrow search with adjusted m/z values. Fragmentation tolerance was set to 700 ppm and 20 mDa and precursor mass tolerance was set to 300 ppm or 10 ppm, respectively. Uniform usage and conversion of parameters into the appropriate command line parameters was done using Ursgal^29^, yielding unified csv as result files. The pyProtista package was used in Ursgal to ensure that the result files for the different engines are comparable and features are uniformly computed. The protein database used as input for all algorithms was Uniprot human and *E. coli* version 2018-01 with all isoleucines converted to leucines to remove ambiguity between algorithms and assignments. The decoy database was generated using ursgal’s unode target_decoy_generation (v2.0) in peptide shuffle mode where each tryptic peptide was shuffled keeping mass, amino acid distribution, and cleavage site constant, thereby keeping the target and decoy search space comparable. Additionally, the target_decoy_generation code ensured that sequences can only exist in the target or decoy space and that non-unique target peptides map onto the same decoy sequence.

### Data

Input data for PeptideForest are:

- Mascot: scores PSMs using the logarithm of the probability *P* that an observed match is a random event, *-10\log10(P)*. A higher score means a better match.
- MSGF+: scores PSMs using the *e*-value. A lower *e*-value means a better match.
- OMSSA: *e*-value.
- MSFragger: scores PSMs using the hyperscore.
- Comet: *e*-value.
- MS Amanda: scoring using the MS Amanda score, a higher score is better

Once the search engine runs are complete and unified using pyProtista, the results are combined into one dataset, and the data is preprocessed to obtain features for training PeptideForest.

### Target-Decoy validation method

These search engines use the target-decoy method to obtain false detection rates (FDR), whereby a database of true (target) peptide spectra is combined with a database of false (decoy) spectra. As any match to a decoy peptide spectrum is known to be a false positive, the ratio of decoy matches with scores below a given score *s’ (M_*{*d*}*(s<s’))* to target matches with scores below a given score *s’ (M_*{*t*}*(s<s’))* normalized by the ratio of target to decoy PSMs in the dataset, *N_*{*t*}*/N_*{*d*}. This is given by the equation below.

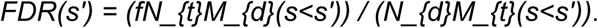

The constant *f* is the estimate of what fraction of the target PSMs are true positives. Since this is unknown, we choose a constant value of *0*.*9* throughout.

### Feature Extraction

The features we use can be placed in three different groups, which we describe below.

- Experimental features: these are experimental measured features from the sample.
  - Mass: the observed mass.
  - Charge_i: if the charge is equal to i. A maximum i value can be set, with any higher values grouped.
- Peptide Features: these are features from the peptide that the spectrum has been matched to. Some are linked to the experimental measurements above.
  - enzN: if the peptide is preceded by an enzymatic (tryptic) site.
  - enzC: if the peptide has an enzymatic (tryptic) C-terminus.
  - enzInt: number of missed internal enzymatic (tryptic) sites.
  - PepLen: length of the sequence of the matched peptide.
  - CountProt: number of proteins the peptide is mapping to.
  - lnNumPep: natural log of the number of PSMs for which this is the best-scoring peptide.
  - DeltaMass: difference between the observed and theoretical masses.
  - matched_b_ions: number of matched b-ions
  - matched_y_ions: number of matched y-ions
  - explained_i_b_ions: percentage of spectrum intensity explained by b-ions
  - explained_i_y_ions: percentage of spectrum intensity explained by y-ions
- Search Engine Features: these are taken from values returned by the search engine.
  - Score_processed: score from the search engine. For Mascot, the score is used as returned by the engine. For MSGF+ and OMSSA, the negative of the log (to base 10) is used.
  - delta_score_i: the difference in Score_processed between a given PSM and the i^{th} ranked PSM for a given experimental spectrum. This feature is only returned if there are at least 70% with multiple PSMs for each spectrum. This was only possible for the MSGF+ and MSFragger search engine results. We limited this to i = 2 and 3.

Features used during training can be dynamically selected during configuration.

### PeptideForest

We build on the approach of the Percolator model, first described in ^30^, which uses a Support Vector Machine (SVM) machine learning model to separate high-quality target PSMs (defined as those with a *q-value <= 1%*) from decoy PSMs. For PeptideForest we use a Random Forest regression (RF-reg) machine learning method to predict if a PSM is a target or a decoy, with the probability that a PSM is a target used as a score, which is used to re-rank PSMs and identify more high-quality target PSMs as matches than the search engines alone. The PeptideForest algorithm can be described as follows:

- Combine results from different search engines
- Calculate features
- Train model:
  1. split data set into three sets at the spectrum level
  2. make a training dataset of two of the splits, 2/3 of all spectra, and a test set from the final split
  3. rank PSMs (by specified engine score during the first iteration and the model score for all subsequent iterations) and take all top-targets (those with *q-value < 1%*) and the same number of randomly chosen decoys from the training dataset
  4. train model to separate targets from decoys on the training set
  5. score all PSMs in the test set using the trained model
  6. repeat steps 2 - 5 twice more, using each combination of training and test sets constructed by the three splits. Each split is scored using a model trained on other data
  7. rerank PSMs by the PeptideForest score
  8. Repeat steps 4 - 7 n_iter times, and average the score over the final n_eval iterations
  9. final scores are the average of the scores obtained from each initial scoring engine
- calculate *q-values* based on final average scores
  1. rank PSMs by scores from the machine learning model
  2. take the highest-scoring PSM for each spectrum
  3. if two (or more) PSMs for a given spectrum are both ranked no. 1 with the same score, then:
    - if both are decoys, pick one at random to keep
    - if at least one is a target and at least one is a decoy, drop both
    - if both are targets, drop both
  4. calculate *q-values* as outlined above
- return target PSMs with q-value <= 1%.

## Acknowledgements

We would like to thank Dr. Gitte Neubauer for sponsoring the kick-off of this project.

## Notes

https://github.com/peptideforest/peptideForest

### Competing Interest Statement

The authors have declared no competing interest.

## References

(1) Perkins, D. N.; Pappin, D. J. C.; Creasy, D. M.; Cottrell, J. S. Probability-Based Protein Identification by Searching Sequence Databases Using Mass Spectrometry Data. ELECTROPHORESIS 1999, 20 (18), 3551–3567. https://doi.org/10.1002/(SICI)1522-2683(19991201)20:18<3551::AID-ELPS3551>3.0.CO;2-2.

(2) Craig, R.; Beavis, R. C. A Method for Reducing the Time Required to Match Protein Sequences with Tandem Mass Spectra. Rapid Commun. Mass Spectrom. 2003, 17 (20), 2310–2316. https://doi.org/10.1002/rcm.1198.

(3) Geer, L. Y.; Markey, S. P.; Kowalak, J. A.; Wagner, L.; Xu, M.; Maynard, D. M.; Yang, X.; Shi, W.; Bryant, S. H. Open Mass Spectrometry Search Algorithm. J. Proteome Res. 2004, 3 (5), 958–964. https://doi.org/10.1021/pr0499491.

(4) Tabb, D. L.; Fernando, C. G.; Chambers, M. C. MyriMatch: Highly Accurate Tandem Mass Spectral Peptide Identification by Multivariate Hypergeometric Analysis. J. Proteome Res. 2007, 6 (2), 654–661. https://doi.org/10.1021/pr0604054.

(5) Park, C. Y.; Klammer, A. A.; Käll, L.; MacCoss, M. J.; Noble, W. S. Rapid and Accurate Peptide Identification from Tandem Mass Spectra. J. Proteome Res. 2008, 7 (7), 3022–3027. https://doi.org/10.1021/pr800127y.

(6) Dorfer, V.; Pichler, P.; Stranzl, T.; Stadlmann, J.; Taus, T.; Winkler, S.; Mechtler, K. MS Amanda, a Universal Identification Algorithm Optimized for High Accuracy Tandem Mass Spectra. J. Proteome Res. 2014, 13 (8), 3679–3684. https://doi.org/10.1021/pr500202e.

(7) Kim, S.; Pevzner, P. A. MS-GF+ Makes Progress towards a Universal Database Search Tool for Proteomics. Nat. Commun. 2014, 5 (1), 5277. https://doi.org/10.1038/ncomms6277.

(8) Kong, A. T.; Leprevost, F. V.; Avtonomov, D. M.; Mellacheruvu, D.; Nesvizhskii, A. I. MSFragger: Ultrafast and Comprehensive Peptide Identification in Mass Spectrometry–Based Proteomics. Nat. Methods 2017, 14 (5), 513–520. https://doi.org/10.1038/nmeth.4256.

(9) Shiferaw, G. A.; Vandermarliere, E.; Hulstaert, N.; Gabriels, R.; Martens, L.; Volders, P.-J. COSS: A Fast and User-Friendly Tool for Spectral Library Searching. J. Proteome Res. 2020, 19 (7), 2786–2793. https://doi.org/10.1021/acs.jproteome.9b00743.

(10) Bruderer, R.; Bernhardt, O. M.; Gandhi, T.; Miladinović, S. M.; Cheng, L.-Y.; Messner, S.; Ehrenberger, T.; Zanotelli, V.; Butscheid, Y.; Escher, C.; Vitek, O.; Rinner, O.; Reiter, L. Extending the Limits of Quantitative Proteome Profiling with Data-Independent Acquisition and Application to Acetaminophen-Treated Three-Dimensional Liver Microtissues. Mol. Cell. Proteomics 2015, 14 (5), 1400–1410. https://doi.org/10.1074/mcp.M114.044305.

(11) Frewen, B. E.; Merrihew, G. E.; Wu, C. C.; Noble, W. S.; MacCoss, M. J. Analysis of Peptide MS/MS Spectra from Large-Scale Proteomics Experiments Using Spectrum Libraries. Anal. Chem. 2006, 78 (16), 5678–5684. https://doi.org/10.1021/ac060279n.

(12) Lam, H.; Deutsch, E. W.; Eddes, J. S.; Eng, J. K.; King, N.; Stein, S. E.; Aebersold, R. Development and Validation of a Spectral Library Searching Method for Peptide Identification from MS/MS. PROTEOMICS 2007, 7 (5), 655–667. https://doi.org/10.1002/pmic.200600625.

(13) Stein, S. E.; Scott, D. R. Optimization and Testing of Mass Spectral Library Search Algorithms for Compound Identification. J. Am. Soc. Mass Spectrom. 1994, 5 (9), 859–866. https://doi.org/10.1016/1044-0305(94)87009-8.

(14) Ma, B. Novor: Real-Time Peptide de Novo Sequencing Software. J. Am. Soc. Mass Spectrom. 2015, 26 (11), 1885–1894. https://doi.org/10.1007/s13361-015-1204-0.

(15) Jeong, K.; Kim, S.; Pevzner, P. A. UniNovo: A Universal Tool for de Novo Peptide Sequencing. Bioinformatics 2013, 29 (16), 1953–1962. https://doi.org/10.1093/bioinformatics/btt338.

(16) Yang, H.; Chi, H.; Zeng, W.-F.; Zhou, W.-J.; He, S.-M. PNovo 3: Precise de Novo Peptide Sequencing Using a Learning-to-Rank Framework. Bioinformatics 2019, 35 (14), i183–i190. https://doi.org/10.1093/bioinformatics/btz366.

(17) Tran, N. H.; Zhang, X.; Xin, L.; Shan, B.; Li, M. De Novo Peptide Sequencing by Deep Learning. Proc. Natl. Acad. Sci. U. S. A. 2017, 114 (31), 8247–8252. https://doi.org/10.1073/pnas.1705691114.

(18) Devabhaktuni, A.; Lin, S.; Zhang, L.; Swaminathan, K.; Gonzales, C.; Olsson, N.; Pearlman, S.; Rawson, K.; Elias, J. E. TagGraph Reveals Vast Protein Modification Landscapes from Large Tandem Mass Spectrometry Data Sets. Nat. Biotechnol. 2019, 37 (4), 469–479. https://doi.org/10.1038/s41587-019-0067-5.

(19) Na, S.; Bandeira, N.; Paek, E. Fast Multi-Blind Modification Search through Tandem Mass Spectrometry. Mol. Cell. Proteomics MCP 2012, 11 (4), M111.010199. https://doi.org/10.1074/mcp.M111.010199.

(20) Yu, F.; Li, N.; Yu, W. PIPI: PTM-Invariant Peptide Identification Using Coding Method. J. Proteome Res. 2016, 15 (12), 4423–4435. https://doi.org/10.1021/acs.jproteome.6b00485.

(21) Levitsky, L. I.; Ivanov, M. V.; Lobas, A. A.; Bubis, J. A.; Tarasova, I. A.; Solovyeva, E. M.; Pridatchenko, M. L.; Gorshkov, M. V. IdentiPy: An Extensible Search Engine for Protein Identification in Shotgun Proteomics. J. Proteome Res. 2018, 17 (7), 2249–2255. https://doi.org/10.1021/acs.jproteome.7b00640.

(22) Solntsev, S. K.; Shortreed, M. R.; Frey, B. L.; Smith, L. M. Enhanced Global Post-Translational Modification Discovery with MetaMorpheus. J. Proteome Res. 2018, 17 (5), 1844–1851. https://doi.org/10.1021/acs.jproteome.7b00873.

(23) Eng, J. K.; McCormack, A. L.; Yates, J. R. An Approach to Correlate Tandem Mass Spectral Data of Peptides with Amino Acid Sequences in a Protein Database. J. Am. Soc. Mass Spectrom. 1994, 5 (11), 976–989. https://doi.org/10.1016/1044-0305(94)80016-2.

(24) Craig, R.; Beavis, R. C. TANDEM: Matching Proteins with Tandem Mass Spectra. Bioinforma. Oxf. Engl. 2004, 20 (9), 1466–1467. https://doi.org/10.1093/bioinformatics/bth092.

(25) Jones, A. R.; Siepen, J. A.; Hubbard, S. J.; Paton, N. W. Improving Sensitivity in Proteome Studies by Analysis of False Discovery Rates for Multiple Search Engines. Proteomics 2009, 9 (5), 1220–1229. https://doi.org/10.1002/pmic.200800473.

(26) Kwon, T.; Choi, H.; Vogel, C.; Nesvizhskii, A. I.; Marcotte, E. M. MSblender: A Probabilistic Approach for Integrating Peptide Identifications from Multiple Database Search Engines. J. Proteome Res. 2011, 10 (7), 2949–2958. https://doi.org/10.1021/pr2002116.

(27) Shteynberg, D.; Nesvizhskii, A. I.; Moritz, R. L.; Deutsch, E. W. Combining Results of Multiple Search Engines in Proteomics. Mol. Cell. Proteomics MCP 2013, 12 (9), 2383–2393. https://doi.org/10.1074/mcp.R113.027797.

(28) Vaudel, M.; Burkhart, J. M.; Zahedi, R. P.; Oveland, E.; Berven, F. S.; Sickmann, A.; Martens, L.; Barsnes, H. PeptideShaker Enables Reanalysis of MS-Derived Proteomics Data Sets. Nat. Biotechnol. 2015, 33 (1), 22–24. https://doi.org/10.1038/nbt.3109.

(29) Kremer, L. P. M.; Leufken, J.; Oyunchimeg, P.; Schulze, S.; Fufezan, C. Ursgal, Universal Python Module Combining Common Bottom-Up Proteomics Tools for Large-Scale Analysis. J. Proteome Res. 2016, 15 (3), 788–794. https://doi.org/10.1021/acs.jproteome.5b00860.

(30) Käll, L.; Canterbury, J. D.; Weston, J.; Noble, W. S.; MacCoss, M. J. Semi-Supervised Learning for Peptide Identification from Shotgun Proteomics Datasets. Nat. Methods 2007, 4 (11), 923–925. https://doi.org/10.1038/nmeth1113.

(31) The, M.; MacCoss, M. J.; Noble, W. S.; Käll, L. Fast and Accurate Protein False Discovery Rates on Large-Scale Proteomics Data Sets with Percolator 3.0. J. Am. Soc. Mass Spectrom. 2016, 27 (11), 1719–1727. https://doi.org/10.1007/s13361-016-1460-7.

(32) Savitski, M. M.; Sweetman, G.; Askenazi, M.; Marto, J. A.; Lang, M.; Zinn, N.; Bantscheff, M. Delayed Fragmentation and Optimized Isolation Width Settings for Improvement of Protein Identification and Accuracy of Isobaric Mass Tag Quantification on Orbitrap-Type Mass Spectrometers. Anal. Chem. 2011, 83 (23), 8959–8967. https://doi.org/10.1021/ac201760x.

(33) Savitski, M. M.; Mathieson, T.; Zinn, N.; Sweetman, G.; Doce, C.; Becher, I.; Pachl, F.; Kuster, B.; Bantscheff, M. Measuring and Managing Ratio Compression for Accurate ITRAQ/TMT Quantification. J. Proteome Res. 2013, 12 (8), 3586–3598. https://doi.org/10.1021/pr400098r.

(34) Smits, A. H.; Ziebell, F.; Joberty, G.; Zinn, N.; Mueller, W. F.; Clauder-Münster, S.; Eberhard, D.; Fälth Savitski, M.; Grandi, P.; Jakob, P.; Michon, A.-M.; Sun, H.; Tessmer, K.; Bürckstümmer, T.; Bantscheff, M.; Steinmetz, L. M.; Drewes, G.; Huber, W. Biological Plasticity Rescues Target Activity in CRISPR Knock Outs. Nat. Methods 2019, 16 (11), 1087–1093. https://doi.org/10.1038/s41592-019-0614-5.

(35) Werner, T.; Becher, I.; Sweetman, G.; Doce, C.; Savitski, M. M.; Bantscheff, M. High-Resolution Enabled TMT 8-Plexing. Anal. Chem. 2012, 84 (16), 7188–7194. https://doi.org/10.1021/ac301553x.

(36) Werner, T.; Sweetman, G.; Savitski, M. F.; Mathieson, T.; Bantscheff, M.; Savitski, M. M. Ion Coalescence of Neutron Encoded TMT 10-Plex Reporter Ions. Anal. Chem. 2014, 86 (7), 3594–3601. https://doi.org/10.1021/ac500140s.

(37) Chambers, M. C.; Maclean, B.; Burke, R.; Amodei, D.; Ruderman, D. L.; Neumann, S.; Gatto, L.; Fischer, B.; Pratt, B.; Egertson, J.; Hoff, K.; Kessner, D.; Tasman, N.; Shulman, N.; Frewen, B.; Baker, T. A.; Brusniak, M.-Y.; Paulse, C.; Creasy, D.; Flashner, L.; Kani, K.; Moulding, C.; Seymour, S. L.; Nuwaysir, L. M.; Lefebvre, B.; Kuhlmann, F.; Roark, J.; Rainer, P.; Detlev, S.; Hemenway, T.; Huhmer, A.; Langridge, J.; Connolly, B.; Chadick, T.; Holly, K.; Eckels, J.; Deutsch, E. W.; Moritz, R. L.; Katz, J. E.; Agus, D. B.; MacCoss, M.; Tabb, D. L.; Mallick, P. A Cross-Platform Toolkit for Mass Spectrometry and Proteomics. Nat. Biotechnol. 2012, 30 (10), 918–920. https://doi.org/10.1038/nbt.2377.

(38) Eng, J.K., Jahan, T.A. and Hoopmann, M.R. Comet: An open-source MS/MS sequence database search tool. Proteomics. 2013, 13: 22–24. https://doi.org/10.1002/pmic.201200439

(39) V. Dorfer et al., MS Amanda, a Universal Identification Algorithm Optimized for High Accuracy Tandem Mass Spectra. J. Proteome Res. 2014, 13 (8), 3679–3684. doi: 10.1021/pr500202e.

